# Ribosome Profiling in Archaea Reveals Leaderless Translation, Novel Translational Initiation Sites, and Ribosome Pausing at Single Codon Resolution

**DOI:** 10.1101/2020.02.04.934349

**Authors:** Diego Rivera Gelsinger, Emma Dallon, Rahul Reddy, Fuad Mohammad, Allen R. Buskirk, Jocelyne DiRuggiero

**Affiliations:** Department of Biology and Department of Earth and Planetary Sciences, the Johns Hopkins University, Baltimore, Maryland, 21218, USA; Department of Molecular Biology and Genetics, the Johns Hopkins University School of Medicine, Baltimore, Maryland, 21218, USA

## Abstract

High-throughput methods, such as ribosome profiling, have revealed the complexity of translation regulation in Bacteria and Eukarya with large-scale effects on cellular functions. In contrast, the translational landscape in Archaea remains mostly unexplored. Here, we developed ribosome profiling in a model archaeon, *Haloferax volcanii*, elucidating, for the first time, the translational landscape of a representative of the third domain of life. We determined the ribosome footprint of *H. volcanii* to be comparable in size to that of the Eukarya. We linked footprint lengths to initiating and elongating states of the ribosome on leadered transcripts, operons, and on leaderless transcripts, the latter representing 70% of *H. volcanii* transcriptome. We manipulated ribosome activity with translation inhibitors to reveal ribosome pausing at specific codons. Lastly, we found that the drug harringtonine arrested ribosomes at initiation sites in this archaeon. This drug treatment allowed us to confirm known translation initiation sites and also reveal putative novel initiation sites in intergenic regions and within genes. Ribosome profiling revealed an uncharacterized complexity of translation in this archaeon with bacteria-like, eukarya-like, and potentially novel translation mechanisms. These mechanisms are likely to be functionally essential and to contribute to an expanded proteome with regulatory roles in gene expression.

## INTRODUCTION

Archaea stand at the crossroad between the other two domains of life, using a mosaic of molecular features from both Bacteria and Eukarya along with unique characteristics. For example, Archaea have a eukaryotic-like transcription apparatus but bacterial-like transcription regulation and coupled transcription-translation (1). Moreover, Archaea have polycistronic mRNAs like Bacteria, implying that archaeal ribosomes can perform repeated cycles of initiation on the same mRNA (2–4). Archaea are prokaryotes and, as such, their ribosome subunits (30S/50S) and rRNA genes (16S/23S) are closer in size to their bacterial counterparts rather than to Eukarya (5). Even with these similarities, Archaea are still evolutionarily closer to Eukarya, and recently it has been proposed that the newly discovered Asgard archaea may be sister taxa to the Eukarya (6–9). This close evolutionary relationship between the two domains is based on similarities of central dogma processes (replication, transcription, and translation) and an expected complexity in the Archaea (10, 11). We thus stand to discover novel mechanisms and processes by studying the molecular biology of Archaea.

Translation is essential for life and has been proposed to be the first subsystem to “crystallize” during the transition of pre-cellular to cellular life as we know it (12). Despite evolutionary conservation, the translation apparatus, the ribosomes, and their accessory proteins have diverged in the three domains of life (11, 13, 14). The bacterial translation system is highly streamlined, while Archaea and Eukarya have evolved an expansion of initiation and recycling factors that causes a high degree of complexity in the translation cycle and suggests a need for extensive regulation of this process (11, 13, 15, 16).

Even after decades of research, our understanding of translation in Archaea remains limited compared to the other domains of life. There is also a puzzling undeniable link between the archaeal and eukaryotic translation apparatus that remains to be addressed. The lack of knowledge in archaeal translation mechanisms is primarily due to the difficulty in cultivating these organisms and because many archaeal model organisms are extremophiles, making it difficult to use classical biochemical and molecular methods. While substantial work has been done to characterize archaeal mechanisms of translation initiation *in vitro* (14, 17–19), these studies were focused on a few select genes as opposed to a global approach. As a result, it remains difficult to predict a unified view of translation processes and their regulation across all mRNAs in an archaeon where many different transcriptional units co-exist (i.e. leadered, leaderless, and operonic transcripts). There is, therefore, a clear need for a high-resolution, genome-wide view of translation in the Archaea.

To address the current knowledge gap in the mechanisms and regulation of translation in Archaea, we developed a simple, robust, and reproducible protocol for ribosome profiling in the model halophilic archaeon *Haloferax volcanii* (*Hv*). *Hv*, a member of the *Halobacteria*, is adapted to high salt and uses a “salt-in” strategy (accumulation of 2-4 M KCl within the cell) to balance the osmotic pressure from its high salt environment (20). We used ribosome profiling to gain a high-resolution view of various aspects of archaeal translation, including the ribosome footprint size of archaea, leaderless initiation, and global ribosomal pausing. Going further, we used harringtonine to stall ribosomes at initiation codons and identified novel and alternative translation initiation sites genome-wide. This work describes novel insights into archaeal translation processes and provides an experimental paradigm for the *in vivo* study of translation at high salt. It also provides a framework for the adaptation of this technique to other Archaea.

## MATERIAL AND METHODS

*H. volcanii culture conditions, harvesting, and cell lysis*. *H. volcanii* DS2 strain H98 (Δ*pyrE2* Δ*hdrB*) (*Hv*) single colonies were picked and grown overnight at 42°C with shaking at 220 rpm (Amerex Gyromax 737) in Hv-YPC medium supplemented with thymidine (50 μg/mL final concentration) (21) until saturation (OD_600_ > 1.0). These cultures were then diluted to OD_600_ 0.02 in fresh media, grown to OD_600_ 0.4 and split evenly into two flasks; one flask was used as a no-treatment control and the other was treated with either 1 mg/mL homo-harringtonine (HHT, Sigma Catalog #SML1091), 20 mM serine hydroxamate (SHX, Sigma Catalog #S4503), or 100 μg/mL anisomycin (ANS, Sigma Catalog #A9789) with final concentrations as indicated in the text. Cells were harvested by centrifugation, filtering, or direct freezing of the culture in liquid nitrogen.

For cells harvested by centrifugation, cultures were immediately centrifuged at 8,600 x g for 3 min at room temperature (RT), the supernatant removed, and the pellets flashed frozen in liquid nitrogen. For cell lysis, the frozen pellets were resuspended in 1 mL of 1x lysis buffer (3.4 M KCl, 500 mM MgCl_2,_ 50 mM CaCl_2_, 1 M Tris pH 7.5) with an additional 100 μg/mL ANS, transferred to a cryo-mill (Spex SamplePrep 6870), and pulverized with 5 cycles (1 min grinding at 5 Hz, 1 min cooling). Lysates were then thawed at RT, centrifuged at 10,000 x g for 5 min, and transferred to a new tube on ice. For cells harvested by filtration, cultures were immediately poured into a filtration unit (90 mm, Sigma WHA1950009) on a 0.25 μm filter. As soon as cells began to accumulate on the filter (for < 1 min), cells were collected using a sterile spatula, flash-frozen in liquid nitrogen, and lysed in a cryo-mill in the same manner as the centrifugation harvesting method.

For direct freezing of cultures in liquid nitrogen, 100 mL of culture were sprayed directly into liquid nitrogen using a serological pipette. The frozen culture formed small pellets that were collected, and 50 g of pellets were weighed to add 1x lysis buffer (3.4 M KCl, 500 mM MgCl_2,_ 50 mM CaCl_2_, 1 M Tris pH 7.5) and 100 μg/mL ANS to prevent ribosome elongation when thawed later on. The pellets were then pulverized in a cryo-mill for 10 cycles due to the larger volume of input (1 min grinding at 10 Hz, 1 min cooling). The lysates were thawed at RT and ribosomes were pelleted over a 60% sucrose cushion (sucrose dissolved in lysis buffer) in an ultracentrifuge with a Ti-70 rotor at 26,4902 x g (60,000 rpm) for 2 hrs at 4°C. Ribosome pellets were resuspended in 200 μL lysis buffer.

### Determination of translation inhibitor concentrations

The following translation inhibitors were tested in *Hv*: ANS, cycloheximide (Sigma Catalog #C1988), HHT, SHX, thiostrepton (Sigma Catalog #T8902), and tetracycline (Sigma Catalog #T8032). These inhibitors were tested either to prevent ribosome elongation after cell lysis or to alter translation in order to validate our ribosome profiling method. Serial dilutions by one order of magnitude of each drug (ANS 1 μg/mL, 10 μg/mL, 100 μg/mL; cycloheximide: 10 μg/mL, 100 μg/mL, 1 mg/mL; HHT: 500 μg/mL, 1 mg/mL; SHX: 0.2 mM, 2 mM, 20 mM; thiostrepton: 5 μg/mL, 50 μg/mL, 500 μg/mL, tetracycline: 5 μg/mL, 50 μg/mL, 500 μg/mL) were administered to *Hv* liquid cultures and incubated, as described above. 100 μL aliquots were removed to measure the optical density of the culture (600 nm) over a 24-48hrs time-period. Concentrations of drugs that completely halted growth were used as the final concentrations in all ribosome profiling libraries.

### Sucrose gradients for ribosomes and subunits

Cell lysates (from cells harvested by centrifugation or direct freezing) were loaded onto a 10-50% sucrose gradient in 1x lysis buffer. Gradients were centrifuged in an ultracentrifuge using a SW41 rotor at 90,140 x g (35,000 rpm) for 2.5 hrs at 4°C. Gradients were then fractionated into 400 μL fractions to resolve ribosome 30S, 50S subunits, monosomes, and polysome fractions. Fractions were flash frozen on dry ice for later RNA isolation.

### RNA extractions and sucrose fraction characterization

RNA was isolated from sucrose gradient fractions that corresponded to ribosome subunits, monosomes, and polysomes. Briefly, 250 μL Trizol LS (Lifesciences) was added to gradient fractions, vortexed, and incubated at RT for 1 min. 150 µL chloroform was added to the samples, vortexed, and centrifuged at 14,000 x g for 10 min at 4 °C. The mixtures were separated into a lower red phenol-chloroform, an interphase, and a colorless upper aqueous phase that was transferred into a new tube. Glycoblue (NEB) and an equal volume of 2-propanol were added to the samples followed by an incubation of 30 min on dry ice. Samples were then centrifuged at 14,000 x g for 30 min at 4 °C to pellet the RNA. RNA pellets were washed twice with 75% ethanol, air dried for 2 min, and resuspended in nuclease-free water. To determine whether sucrose gradient peaks corresponded to 30S, 50S, monosomes, or polysomes, RNAs extracted from each fraction were separated on a denaturing agarose gel. 16S and 23S rRNA bands were visualized by SYBR Gold staining (ThermoFisher).

### Optimization of MNase activity in high salt buffer

MNase (Nuclease S7, Roche Cat: 10107921001) activity was tested at increasing KCl, MgCl_2_, and CaCl_2_ concentrations in lysis buffer in a 2-step double stranded DNA (dsDNA) absorbance assay. In the first step, serial dilutions of MNase were added to a substrate buffer (2 g/mL salmon sperm dsDNA, 10 mM Tris pH 7.5, lysis buffer) in 96-well plates, shaken for 10 s, and OD at 260 nm was measured every min for 1 h at 25°C. In a second step, the same assay was performed at the optimal concentration of MNase with increasing concentrations of CaCl_2_ in the lysis buffer.

### Footprinting of ribosomes and Ribosome profiling library preparation

Cell lysates (from cells harvested by centrifugation or direct freezing) were processed by first treating 20 absorbance units (AU, Nanodrop) of lysate RNA with 12,000 units of MNase for 1 hr at 25°C. After nuclease digestion, samples were loaded onto 10-50% sucrose gradients and RNA was isolated from monosome fractions as described above. Library preparation was performed as previously described (22). Briefly, 10 µg of RNA fragments were used to purify 10-45 nt RNA fragments by gel electrophoresis (PAGE 15% TBE 7 M Urea gel); RNA fragments were treated with T4 polynucleotide kinase (NEB), ligated to the linker (NEB Universal miRNA Cloning Linker) using T4 RNA ligase (NEB), and gel extracted from a 10% TBE Urea gel. rRNAs were subtracted from the RNA fragments using the Ribo-Zero rRNA removal kit for bacteria (Illumina). The resulting fragments were reverse transcribed with SuperScript III (Invitrogen) using custom primers previously described (22). Template RNA was degraded using NaOH for 20 min at 98 °C, the fragment were gel extracted from a 10% TBE Urea gel, circularized using CircLigase (Epicentre) and PCR amplified (8-12 cycles) with Phusion polymerase (NEB) using custom primers (22). PCR products were gel extracted from a 10% TBE gels and analyzed for size and concentration using a BioAnalyzer high sensitivity DNA kit (standard protocol) before sequencing on an Illumina HiSeq 2500 at the Johns Hopkins Genetic Resources Core Facility (Baltimore, MD).

### Transcriptome reannotation

We used RNA-seq data we previously published (23) and the program Rockhopper2 (default settings for paired-end reads) (24) to accurately determine coordinates for all transcription start sites in the *Hv* transcriptome. Using the output of Rockhopper2, an annotation file (.gff) was created to account for untranslated regions (UTR) both 5’ and 3’ with respect to previously annotated translation start sites (TSS) as well as operonic transcripts. All subsequent ribosome profiling analysis used these transcriptome reannotations instead of database deposited annotations (e.g., NCBI and UCSC).

### Ribosome profiling data analysis

All ribosome profiling data were analyzed using previously established methods and python scripts in *E. coli* (22). In brief, reads were trimmed using trim_galore, reads corresponding to rRNA and tRNA were discarded, and the remaining reads were mapped against the *Hv* NCBI RefSeq genome (taxonomy identification [taxid] 2246), allowing two mismatches using Bowtie v 0.12.7 (25). These alignments were then used in custom python scripts to: (a) calculate read density (assigned to the 3’-end of reads) across the genome, (b) calculate average ribosome position and read length distributions along ORFs using meta-gene analysis (3’- or 5’-end of reads as noted) on all transcripts, leadered transcripts, or leaderless transcripts, (c) calculate average amino acid and codon pause scores, (d) asymmetry scores for ORFs, and (e) expression for each gene (22). Custom bash/awk scripts were used to separate 17 nt and 27 nt footprint reads and these alignments were analyzed in the same manner.

TSS identification with harringtonine (HHT) ribosome profiling libraries was done using previously published custom python scripts that were modified for our purposes (26, 27). Identification of novel TSS and smORFs was done as previously published (26, 27) with the following changes: (a) rpkm was calculated for each TSS peak in both the HHT and no challenge library, (b) the TSS peak rpkm were >5-fold enriched in HHT compared to no challenge, and (c) the ORF rpkm were >5-fold enriched in no challenge compared to HHT. Identification of known TSS, internal TSS (iTSS), and N-terminal extensions was done as previously published using density files assigned to the 3’-end of reads and adjusted for the P-site (15 nt offset) (26).

Translation efficiency analysis was done using previously published RNA-seq data (23). Thresholding of expressed CDS were done using custom python scripts, and transcript per million (TPM) was calculated for each CDS and aTSS using ribosome profiling and corresponding RNA-seq samples. Translation efficiency was calculated by dividing the ribosome profiling TPM by the RNA-seq TPM per CDS and per aTSS.

### tRNA northern blot of tRNA charging

To determine tRNA charging of serine and methionine tRNAs, a tRNA deacylation and β-elimination treatment was performed as previously described (22). Briefly, total RNA was extracted from no treatment and SHX-treated flash frozen cells using the Quick RNA extraction kit (Zymo) following the standard protocol. The RNA was then DNase I (NEB) treated for two hrs using 2 units of DNase I per hour and was cleaned of residual DNase I with the RNA clean and concentrator-5 (Zymo). DNase-treated RNA was then divided into two equal aliquots; one of the aliquots was deacylated by treatment with 1M Tris pH 9 at 37 °C for 1 hr, and then ethanol precipitated. Following deacylation, both aliquots were treated with sodium periodate and 1M lysine to promote β-elimination of oxidized 3’ RNA ends, and then ethanol precipitated. Samples were then run on a 10% TBE 7 M Urea denaturing polyacrylamide gel. RNA was transferred using a wet transfer apparatus (Hoefer TE62) onto a nylon membrane and UV crosslinked to the membrane using the automatic setting (UV Stratalinker 1800). Membranes were probed in Ultrahyb Oligo buffer (Ambion) with 5’-32P-labeled (tRNA^Ser^) TCACGTGTCCGAATGGACAGTAGA or 5’-32P-labeled (tRNA^Met^) ATGAGCCCGGCGGAATCTCCT and signal was detected on a Typhoon phosphoimager.

## RESULTS

### Development of a ribosome profiling method for Haloferax volcanii

Ribosome profiling is the deep sequencing of ribosome-protected mRNA footprints (RPFs) after nuclease digestion. Developing this method for *Hv* provided, for the first time, a genome-wide survey of translation at a level of resolution currently unknown in the Archaea. The method has four critical steps (Fig. 1A): (i) inhibit translation so that ribosomes are arrested as samples are processed, (ii) digest unprotected mRNA with nucleases, (iii) purify 70S monosomes from sucrose gradients, and (iv) convert RPFs to dsDNA for deep sequencing (22). Ensuring that these steps are reproducible and robust in *Hv* was particularly challenging due to its extremely high intracellular salt concentration (2-4 M KCl) and a proteome adapted to these high salt conditions.

**FIGURE 1:**
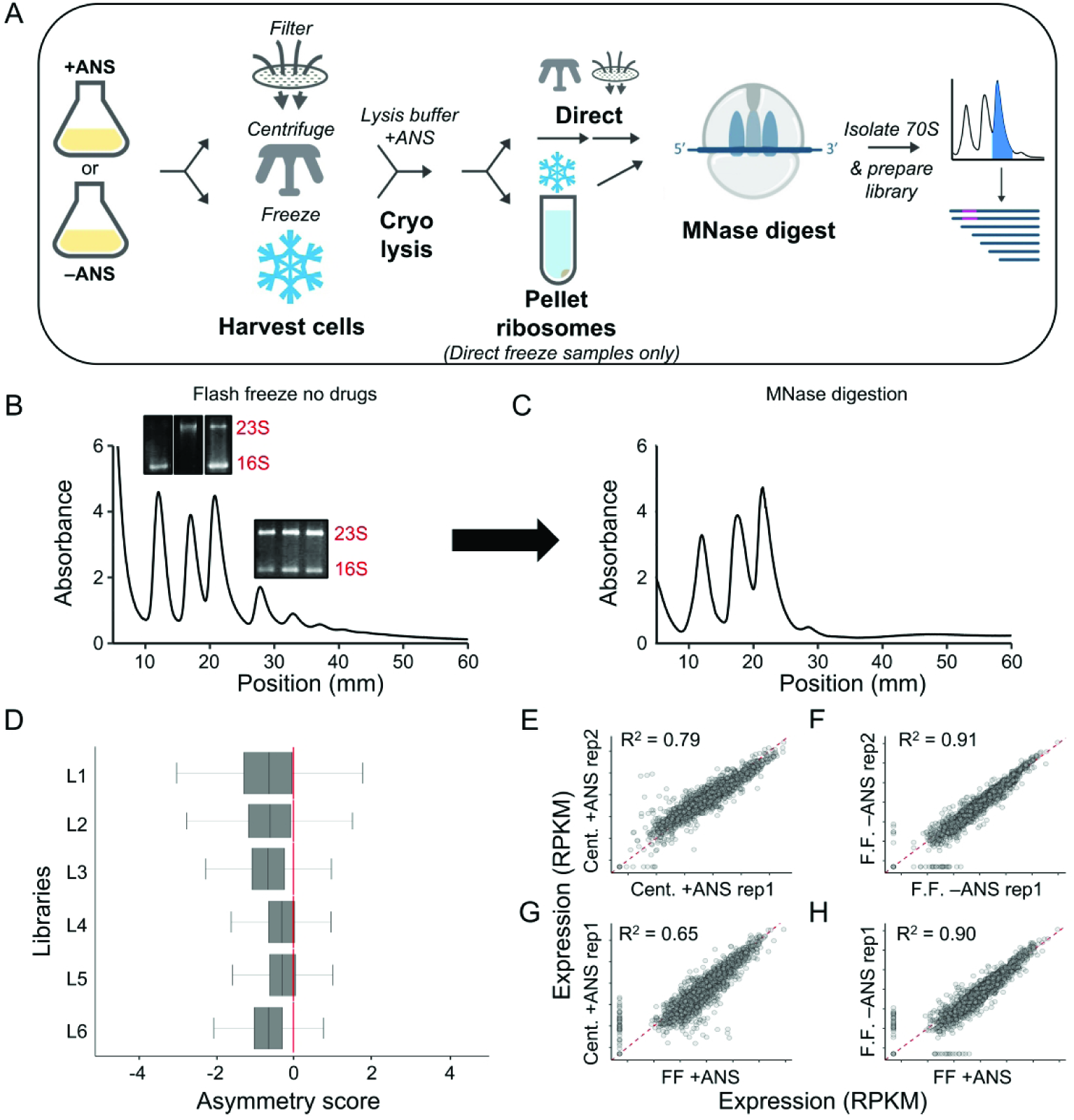
Evaluation of ribosome profiling protocols for *Hv*. (A) Schematic of the ribosome profiling methods used in this study. Sucrose gradients of ribosomes from cells harvested by direct flash freezing in liquid nitrogen with no drugs added to cultures showing (A) intact monosome and polysome fractions, and (B) successful collapse of polysomes into monosomes after MNase digestion. Inset: Total RNA agarose gels showing rRNA from various peaks corresponding to 30S, 50S, 70S, and polysome fractions. (C) Asymmetry scores were calculated as the normalized ribosome density upstream of the halfway point of an ORF over the ribosome density downstream of the halfway point of an ORF. L1= Cent + ANS rep1, L2= Cent + ANS rep2, L3= FF + ANS, L4= FF no drug rep. 1, L5= FF no drug rep. 2, L6= FF + Harringtonine). (D-G) Correlation plots of translation expression (in rpkm) for ribosome profiling libraries of various conditions. ANS, anisomycin; Cent, centrifugation; FF, flash-freezing; rep, biological replicate.

In ribosome profiling studies in bacteria and yeast, cultures are commonly harvested by either filtration or centrifugation. After filtration (for < 1 min), cells were collected from the filter, flash-frozen in liquid nitrogen, and lysed in a cryo-mill. In *Hv*, the filtration procedure did not yield intact ribosomes (Fig. S1A). Using centrifugation at room temperature for 3 min, we observed 30S/50S subunits but very few translating ribosomes (70S monosomes and polysomes, Fig. S1B). This result suggests that *Hv* ribosomes ran off messages during centrifugation, perhaps due to the drop in temperature from 42 °C to 25 °C (29). To arrest ribosomes prior to centrifugation, we tested two antibiotics known to inhibit translational elongation, anisomycin and cycloheximide. We found that adding anisomycin (ANS) to the culture prior to centrifugation halted growth and stabilized translating ribosomes during the centrifugation step (Fig. S1C & Fig. S2). Centrifugation in the presence of ANS, therefore, offered one potential solution for harvesting *Hv* cultures for ribosome profiling.

We also tested a third method for harvesting cells that we recently developed for *E. coli* (22). In this method, cultures were sprayed directly into liquid nitrogen. Flash freezing the cells in the media arrests translation immediately without the use of antibiotics. The frozen cultures were then lysed in a cryo-mill and, in an extra step, ribosomes were pelleted and resuspended into a high-salt lysis buffer (3.4 M KCl, 500 mM MgCl_2_, 10 mM Tris-HCl pH 7.5) with 100 μg/mL ANS to inhibit translation in the lysate. We found that this flash-freezing method yielded the highest level of translating ribosomes (70S monosomes and polysomes) relative to free 30S and 50S subunits (compare Fig. 1B with Fig. S1C). These data show that the flash-freezing protocol developed for bacteria yielded robust polysomes from *Hv* cultures and indicated that ribosomes were arrested and remained intact throughout the purification procedure.

Due to high KCl concentration in the lysis buffer, RNA digestion with micrococcal nuclease (MNase) was inefficient and lowering KCl concentrations to 1 M or 2 M destabilized 70S ribosomes and favored subunit splitting (Fig. S3). Reasoning that high concentrations of KCl might prevent the catalytic metal ion Ca^2+^ from binding in the active site of MNase, we titrated CaCl_2_ in the RNA digestion reaction. We found that with 50 mM CaCl_2_ in the lysis buffer, MNase was able to collapse polysomes into monosomes (Fig. 1C). After purifying 70S monosomes from a sucrose gradient, we isolated 10 – 45 nt mRNA fragments and prepared cDNA libraries for deep sequencing.

Prior studies in yeast and *E. coli* have shown that adding antibiotics to the media introduces artifacts in ribosome profiling studies by trapping ribosomes near the 5’-end of ORFs as elongation is blocked while initiation continues (22, 30). To ask whether the same problem occurs in *Hv*, we sequenced libraries prepared by centrifugation with ANS pre-treatment (Fig. 1D, L1 and L2) and by flash-freezing cultures with (L3) or without (L4 and L5) ANS pre-treatment. We then computed asymmetry scores for each gene, taking the ratio of ribosome density in the second half of the gene over the density in the first half (log_2_ transformed). If the asymmetry score is less than zero, ribosome density is accumulating at the 5’-end of the ORF. We found that cultures pre-treated with ANS prior to harvesting (Fig. 1D, L1-3) had significantly lower asymmetry scores than those not treated with ANS (L4 and L5), regardless of the cell harvesting method, centrifugation or flash-freezing. These results indicate that adding antibiotics to the culture drastically altered the ribosome position along ORFs.

We further investigated the effects of ANS treatment by determining the relative level of gene translation (reported in reads per kilobase per million mapped reads, or rpkm) in libraries of biological replicates prepared by centrifugation and by flash-freezing. Translational levels for each gene were more highly correlated between the two replicates harvested by flash freezing (R^2^ = 0.91, Fig. 1F) than between those harvested by centrifugation (R^2^ = 0.79, Fig. 1E), indicating that flash freezing is a more reproducible harvesting method. This correlation was reduced when comparing centrifugation and flash-freezing methods in cultures pre-treated with ANS (R^2^ < 0.65, Fig. 1G). In contrast, there was a strong correlation between flash freezing samples, even when one culture was treated with ANS and the other was not (R^2^=0.90, Fig. 1H), indicating that the harvesting method (i.e. centrifugation) is the critical difference that alters translational levels and that ANS treatment is less of a factor. Ultimately, we concluded that flash freezing without ANS pre-treatment introduced the fewest artifacts and thus all subsequent libraries described below were prepared with this method.

### 27 nt RPFs reveal the reading frame of the ribosome in Hv

Isolating RPFs between 10 – 45 nt, we consistently found in our ribosome profiling libraries that the largest number of RPFs were 27 nt (Fig. 2A). The distribution of RPFs lengths was reproducible, with only subtle differences between libraries regardless of how they were prepared (by flash freezing or centrifugation, with or without ANS pre-treatment). The predominant 27 nt footprints in *Hv* (Fig. 2B, green) were only slightly shorter than the predominant footprints in eukaryotes (28 – 30 nt, red, orange, and brown). In contrast, most RPFs in *E. coli* are shorter, around 24 nt (blue). In addition, we observed that the distributions for all these species have more than one peak. In yeast and mammalian cells, there are two distributions of RPF lengths, one centered at 28 nt and the other at 21 nt. These two peaks represent two different conformational states of elongating ribosomes (31, 32). In all of our *Hv* ribosome profiling libraries, we also observe a peak of short RPFs (Fig. 2B), but as described below, they differ from the short reads in eukaryotes.

**FIGURE 2:**
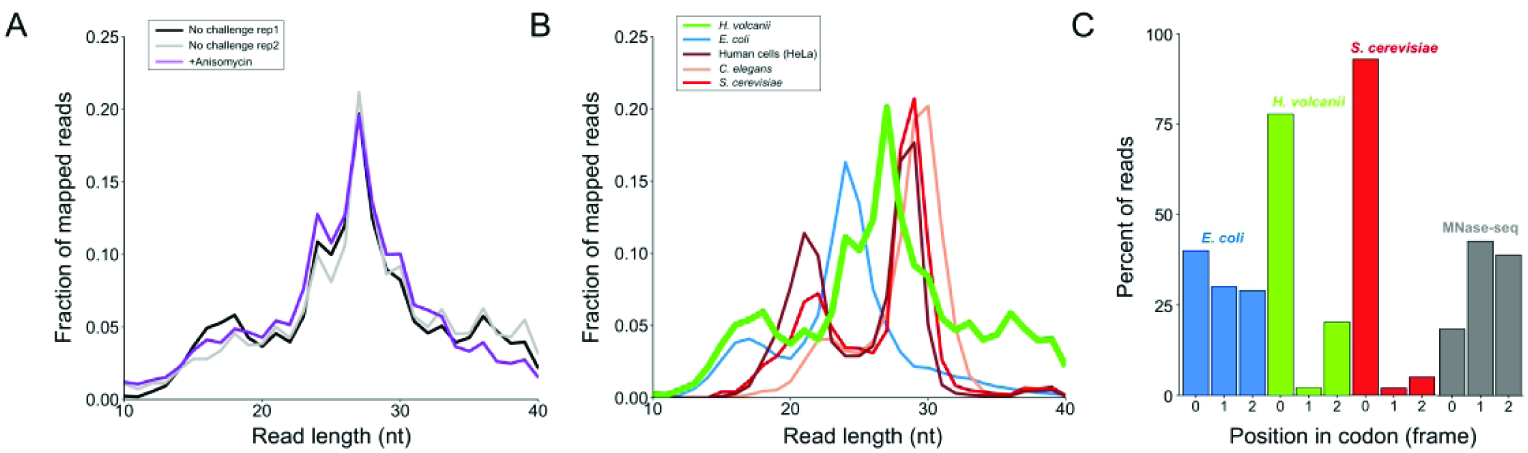
Length distribution of ribosome footprints in *Hv.* (A) Fraction of reads of various lengths in *Hv* ribosome profiling libraries from cultures harvested by flash-freezing without (2 biological replicates) and with anisomycin (ANS) treatment prior to harvesting. (B) Fraction of reads of various lengths in ribosome profiling libraries for *E. coli, S. cerevisiae*, *C. elegans*, mammalian cells (HeLa), and *Hv* cells. (C) Mapped distribution of the elongating footprints to the position in codon (0, +1, or +2) for *E. coli*, *Hv*, *S. cerevisiae*, and total *Hv* RNA treated with MNase (MNase-seq).

Footprint length is one of the hallmarks of active translation that is captured by ribosome profiling: elongation occurs one codon at a time. Because 28 nt RPFs in yeast are fully trimmed by nucleases back to the 5’- and 3’-boundaries of the ribosome, they reveal the position of the ribosome on a mRNA at higher resolution than RPFs of other lengths. 90% of 28 nt RPFs in yeast map to the first nt in codons, evidence of strong three nt periodicity (Fig. 2C, red). Likewise, we observe in *Hv* that the majority of 27 nt RFPs (> 75%, green) map to the first nt of codons (Fig. 2C, green), demonstrating that these RFPs arise from elongating ribosomes moving one codon at a time along mRNAs. These results contrast sharply with those in bacteria: the periodicity in *E. coli* is much weaker with a nearly equal distribution of mapped footprints to the first (40%), second (30%), and third nt (30%) of codons (Fig. 2C, blue). We know that MNase can yield tight distributions of footprints and strong periodicity in yeast ribosome profiling studies (33). To confirm that the three nt periodicity in *Hv* was due to ribosome’s reading frame, we performed RNA-seq using MNase to fragment total RNA, followed by the same steps for library preparation as in ribosome profiling. In the absence of ribosomes, this RNA-seq protocol with MNase yielded a relatively equal distribution of footprints at all three positions of codons (Fig. 2C, gray). The sharp contrast between the lack of periodicity in this RNA-seq protocol and the strong periodicity seen in 27 nt RPFs from ribosome profiling validates that we are accurately following translation at high resolution in *Hv*.

### Hallmarks of actively initiating and elongating ribosomes identified in Hv

The strength of ribosome profiling lies in the averaging of ribosome density across thousands of genes, aligned at their start codons, to perform “meta-gene” analyses. The position of the ribosome on the mRNA can be assigned using either the 5’-end of the footprint, as is commonly done in eukaryotes, or the 3’-end of the footprint, giving better resolution in bacteria (22, 34). Representative examples of meta-gene plots for *Hv* are shown in Fig. 3, using ribosome density with positions assigned either at the 5’-end (Fig. 3A) or 3’-end of footprints (Fig. 3B). These meta-gene plots revealed high ribosome density and three-nucleotide periodicity within ORFs, and lower ribosome density and no periodicity in untranslated regions (Fig. 3A, B). Heatmaps of average ribosome density, separated by footprint size, for both 5’- and 3’-mapped data demonstrated a tight density of 27 nt footprints along ORFs with strong periodicity relative to all other footprint lengths (Fig. 3A, B), consistent with the reading frame analysis shown in Fig. 2C. These findings are consistent with hallmarks of elongating ribosomes observed in previous ribosome profiling studies.

**FIGURE 3:**
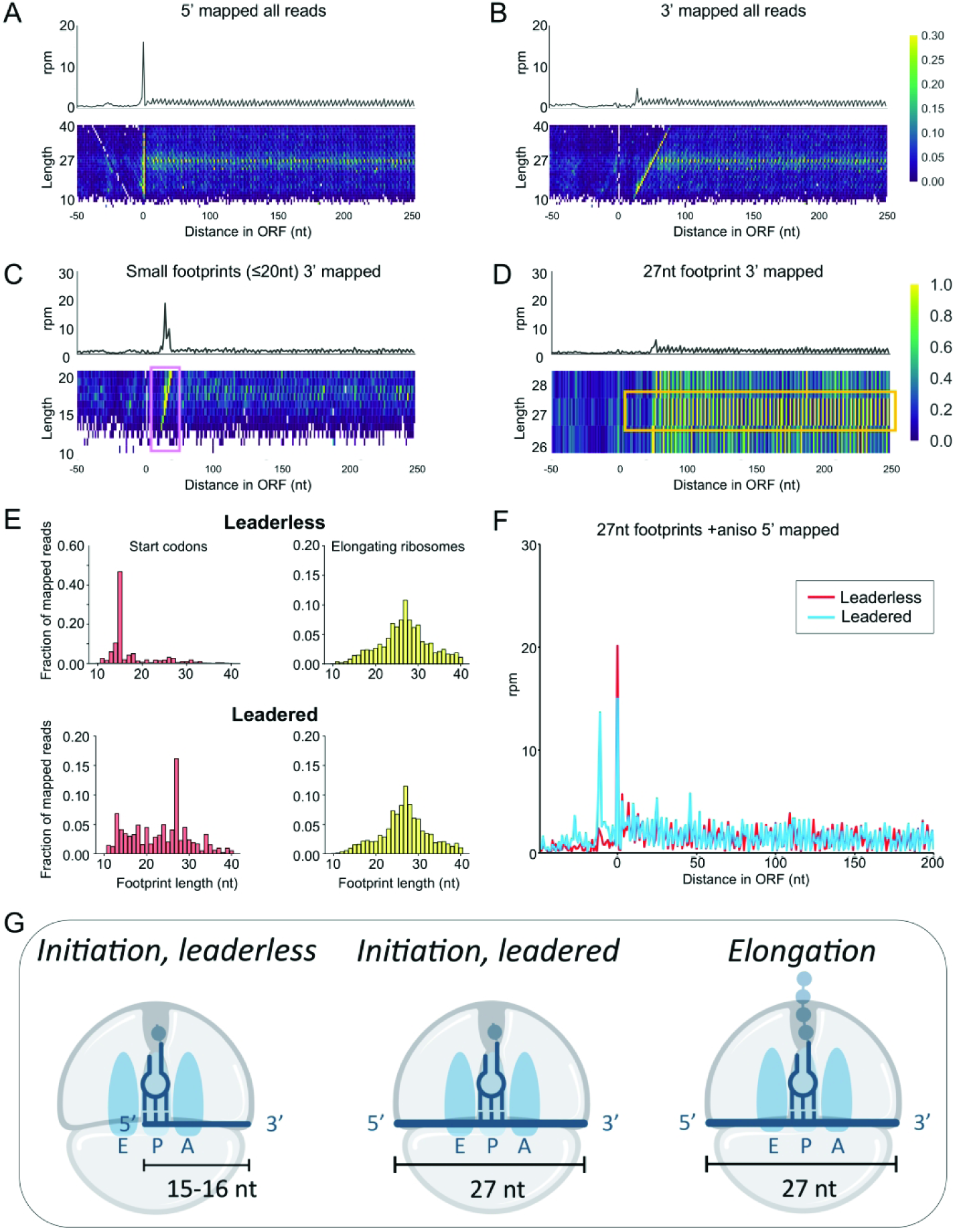
Different length ribosome footprints represent different translational states. Meta-gene plots, from flash frozen and no drug exposure *Hv* cultures, of ribosome density mapped using the (A) 5’-end and (B) 3’-end of ribosome footprints (rpm = reads per million). Heatmaps of footprint density separated by length are aligned underneath the meta-gene plots. (C) 3’-mapped meta-gene analysis of small footprints (10-20 nt) and (D) of 27 nt footprints from flash frozen and no drug cultures. (E) Footprint length distribution of 5’-mapped ribosome density for initiating ribosomes at the start codon (at P-site) in leadered (5’ UTR ≥ 10 nt) and leaderless (0 nt) transcripts (left, red) and elongating ribosomes within the open reading frames (right, yellow). (F) 27 nt footprints on leadered (5’ UTR ≥ 10 nt) and leaderless (0 nt) messenger RNAs from flash frozen and anisomycin (ANS) treatment prior to harvesting. (G) A model for footprint lengths corresponding to ribosome translation states. Small footprints (∼15-16 nt) correspond to initiation on leaderless transcripts, and 27 nt footprints correspond to initiation on leadered transcripts and to elongation on all transcripts.

Another common feature of meta-gene plots is a peak of ribosome density at the start codon that reflects a delay of newly initiated 70S ribosomes in entering the elongation cycle. Although we observed a strong peak in 5’-mapped data (Fig. 3A), the 5’-ends of RPFs were enriched at zero, the first nt in the AUG start codon, rather than enriched at 12 nt upstream of the start codon (the distance between the 5’-boundary of the ribosome and a start codon bound in the P site). Correspondingly, in a heatmap of average ribosome density broken down by footprint size, there was a vertical line of high density at this same position, indicating that the 5’-end of footprints line up with the A in the AUG start codon regardless of the footprint length. In contrast, in the 3’-mapped data (Fig. 3B), the peak near the start codon is significantly lower and shifted downstream. In the associated heatmap broken down by footprint length, there was a diagonal line of high density because the 3’-end of footprints moves downstream as a function of footprint length. Regardless of how we plotted these data, we consistently found an enrichment of footprints whose 5’-ends lies at the first nt of the start codon. We investigated the difference between these footprint sizes further by plotting separately the distribution of small footprints (10-20 nt) and long footprints (26-28 nt) along ORFs using 3’-mapped meta-gene analysis (Fig. 3C-D). The distribution of small footprints showed a bias towards the translation start site, with a peak density at 16 nt, and limited periodicity along ORFs for *Hv* cells with no drug treatment (Fig 3C). In the longer footprint distribution with no drug treatment, the TSS bias was absent and periodicity along the ORF continued downstream, with a peak density at 27 nt (Fig. 3D).

More than 70% of *Hv* mRNAs have been reported to be leaderless (35), starting with the AUG start codon. We reannotated the *Hv* transcriptome with previously published RNA-seq data (23), confirming that the majority of transcripts in *Hv* were indeed leaderless (Fig. S4A). As expected, we observed far fewer reads in RNA-seq data in the 5’-UTR of leaderless transcripts than in leadered transcripts (Fig. S4B). As a result, the enrichment of RPFs at the AUG in Fig. 3A likely arises from the fact that these fragments already have one defined end (the 5’-end). In contrast, fragments in the middle of an mRNA have to be cleaved on both ends to be isolated within the 10–45 nt cutoff and included in the library (36). We conclude that the strong peak in Fig. 3A and 3C at least partially reflects an artifact of sequencing a transcriptome high in leaderless transcripts.

Next, we asked how ribosome footprints differ at start sites on leadered vs leaderless transcripts in *Hv*. As shown in Fig. 3G, we hypothesized that newly-initiated ribosomes on leadered transcripts would protect longer footprints (27 nt) than newly-initiated ribosomes on leaderless transcripts. If the 5’-end of the transcript (the A in AUG) lies in the ribosomal P site, the ribosome should only protect 15-16 nt from nuclease digestion (the distance from the P-site codon to the 3’-boundary of the ribosome). To test this hypothesis, we used the data from L3 (cells treated with ANS) where we saw the most ribosome density at start codons (Fig. S5). For RPFs with start codons in the P site on leaderless transcripts, we observed a strong enrichment of 16 nt RPFs and very few 27 nt RPFs, as expected (Fig. 3E, Leaderless). In contrast, in leadered transcripts, there were significantly more 27 nt RPFs at start codons (Fig. 3E, Leadered) because the ribosome protects additional mRNA upstream of the start codon (as shown in Fig. 3G). In both leadered and leaderless transcripts, RPFs mapping within the ORF tended to be long, with a peak at 27 nt, corresponding to elongating ribosomes.

Another way to visualize these differences at start codons is to plot average ribosome density for only leadered or leaderless genes (Fig. 3F). Using exclusively 27 nt RPFs, we found that their 5’-ends lie at the TSS (zero) in leaderless mRNAs, consistent with two possibilities: (i) the artefactual cloning bias discussed above, and (ii) footprints from newly-initiated ribosomes with no mRNA upstream of the start codon. We cannot distinguish between these two possibilities on leaderless transcripts, but it is likely that both contributed to the signal, because we also observed these on leadered transcripts, where we can differentiate between them. In leadered transcripts, the cloning bias also yields a peak at the TSS (just like leaderless transcripts). But importantly, there is also a strong peak 12 nt upstream of the TSS, consistent with the 5’-end of RPFs with the start codon positioned in the P site (Fig. 3F). This peak corresponds to newly initiated ribosomes.

In summary, we found that elongating ribosomes in *Hv* protect a 27 nt footprint on both leadered and leaderless mRNAs (Fig. 3G). More importantly, our analysis indicated that short footprints (< 20 nt) were strongly enriched at the TSS of leaderless transcripts, either by 5’-cloning artifacts or by bona fide initiation, whereas the 27 nt footprint peak upstream of start codons on leadered mRNAs could be attributed to initiating ribosomes. These observations provide a way to use newly-initiated ribosomes to identify novel initiation sites and better annotate the *Hv* genome, as described below.

### Ribosome profiling detects codon specific translational pauses in Hv

Ribosome profiling studies are able to capture ribosome pausing because when a specific codon is translated more slowly on average, the increased ribosome occupancy indicates that there are more RPFs associated with that codon genome-wide. To test if ribosome pausing could be detected at high resolution in *Hv*, we treated cells with a compound designed to induce pauses at specific codons. Serine hydroxamate (SHX) is an amino acid analog that acts as a competitive inhibitor of seryl-tRNA synthetase in *E. coli* (22, 37). We found that 6 mM SHX halted the growth of *Hv*, indicating that the inhibitor could disrupt its metabolism (Fig. S2). Using ribosome profiling data from cells treated with SHX, we computed pause scores for all 61 sense codons by dividing the ribosome density of the codon of interest by the average density across the gene. When codons were positioned in the E site or the P site of the ribosome, we observed only small deviations from the expected value of 1, indicating little or no pausing (Fig. 4A).

**FIGURE 4:**
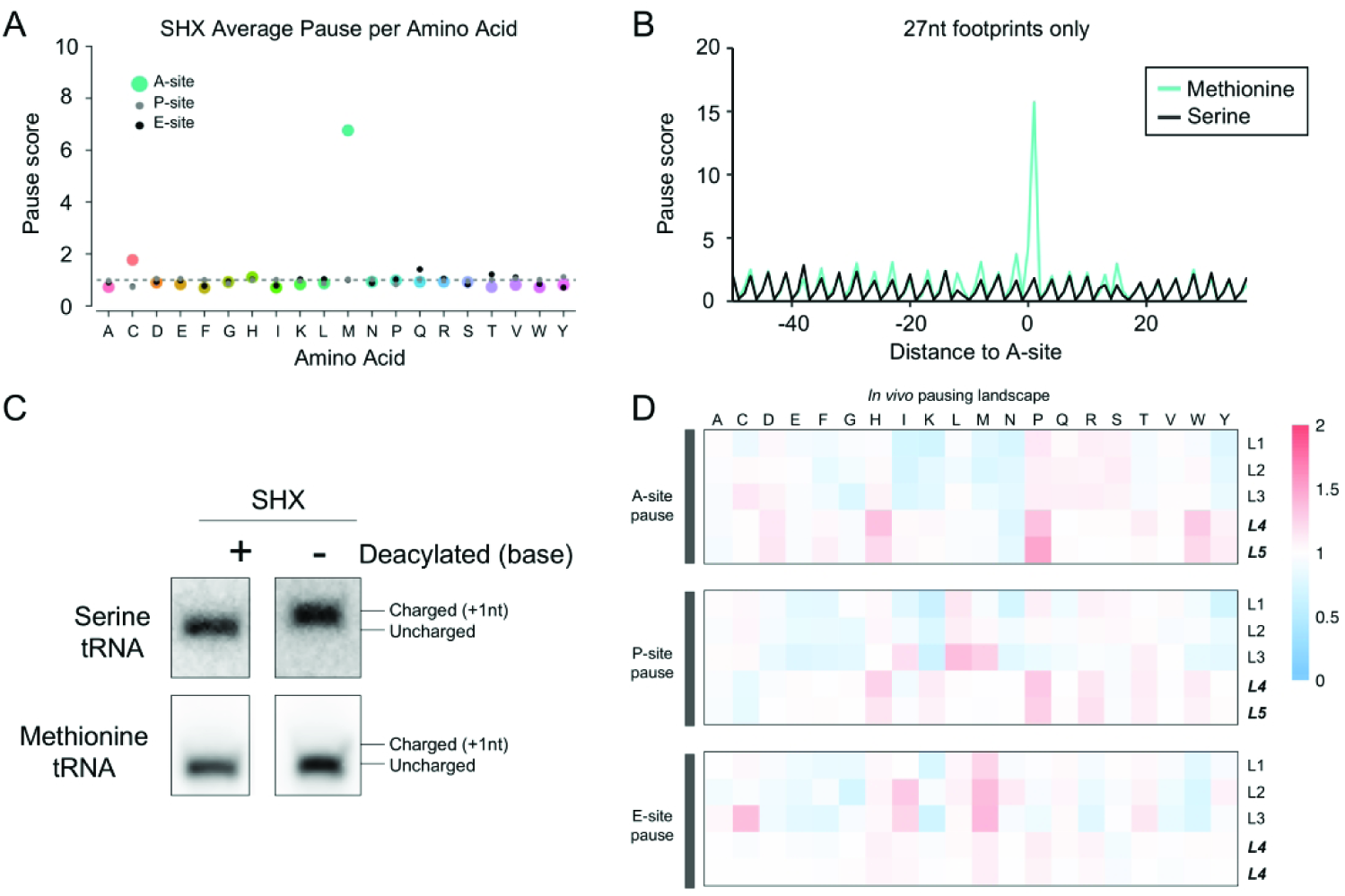
*H. volcanii* global ribosome pausing. (A) Pause scores for all footprints were calculated for each codon and averaged per amino acid at the A-(colored large dot), P-(small gray dot), and E-(small black dot) sites of ribosomes for serine hydroxamate (SHX) treated cells. Ribosome sites were calculated from the 3’ of elongating footprints. (B) Meta-gene plots of ribosome density for elongating footprints (27 nt) for methionine and serine codons. (C) Northern blots of charged and uncharged Met or Ser tRNAs from total RNA of SHX-treated cells. Deacylation treatment is indicated by + and charged/uncharged tRNAs are marked. (D) Comparative heatmap of pause scores for all amino acids under several conditions for A-site, P-site, and E-site of the ribosome.

For the ribosomal A site, however, where decoding takes place, we observed very strong pauses (score = 7) on the codon AUG, which encodes methionine (these calculations did not include start codons). There was also a weaker pause at cysteine codons in the A site (score = 2). These observations are consistent with pausing during decoding as the ribosome waits for a cognate Met or Cys aminoacyl-tRNA that is in low concentration. Surprisingly, there was no evidence of the expected pauses at Ser codons. Limiting our analysis to the 27 nt RPFs revealed the contrast between the Met and Ser codons even more clearly in plots of average ribosome density centered on the codon of interest (Fig. 4B). This additional analysis confirmed that the methionine pauses were specific to the A site of ribosomes and confirmed the lack of pauses at or around Ser codons.

The unexpected observation of strong Met pauses in *Hv* led us to ask if SHX lowers the concentration of Met-tRNA rather than Ser-tRNA as it does in *E. coli*. We extracted total tRNA from *Hv* treated with SHX and, as a control, pre-treated one aliquot with mild base to deacylate all the tRNAs. We then used periodate oxidation and β-elimination to distinguish between charged and uncharged tRNA in these samples. Because uncharged tRNAs are selectively oxidized by periodate, they are one nt shorter than charged tRNAs, allowing their resolution by PAGE and northern blotting using tRNA-specific probes. We observed that in SHX-treated cells, Ser-tRNA (AGC) was fully charged and therefore longer than the deacylated control (Fig. 4C). In contrast, Met-tRNA was uncharged and, therefore, ran at the same size as the deacylated control (Fig. 4C). These results indicate that SHX reduced the level of charged Met-tRNA in *Hv* and not Ser-tRNA, consistent with the A-site pauses observed in the ribosome profiling data. In addition, we found that genes involved in KEGG pathways related to the biosynthesis of methionine and cysteine were up-regulated in ribosome profiling data, reflecting higher levels of expression during SHX treatment; in contrast, serine biosynthesis was unaffected. At this point it remains unclear whether charging of tRNA^Met^ is inhibited directly by SHX or if it acts by blocking the biosynthesis of Met which typically involves Ser as a methyl donor.

Our observations of pauses induced by SHX gave us confidence that we could assess the *in vivo* pausing landscape in *Hv* cells with and without ANS pre-treatment during harvesting. Heatmaps reflecting pause scores for codons in the A-, P-, and E-sites of elongating ribosomes are shown in Fig. 4D. In untreated cells (L4 and L5), the strongest pauses at the A and P sites occurred at Pro codons, consistent with the known biochemistry of Pro as both a poor peptidyl donor and acceptor in the active site of the ribosome. Comparison of the untreated and ANS treated samples confirmed what was previously shown in yeast and in *E. coli* (22, 39, 40), that pre-treatment of cultures with elongation inhibitors (L1-L3) obscures the natural pausing landscape observed in L4 and L5. These data once again highlight the importance of harvesting cells without pre-treatment with antibiotics.

### High-throughput identification of translation start sites

The translation inhibitors harringtonine (HHT) and lactimidomycin have been used in eukaryotic cells to trap ribosomes at initiation sites by inhibiting elongation during the first rounds of peptide-bond formation after subunit joining (28, 41). Importantly, elongating ribosomes are not inhibited by these antibiotics, allowing them to continue elongation and terminate normally at a stop codon (28, 41–43). We reasoned that we could identify translational start sites globally in *Hv* by leveraging the effects of HHT. We determined that 1 mg/mL HHT prevented growth in *Hv* (Fig. S2), a concentration that is an order of magnitude higher than required for eukaryotes. We then conducted ribosome profiling with cells treated with HHT. In meta-gene plots aligned at start codons, we found that ribosomes strongly accumulated at annotated TSS in the presence of HHT compared with untreated samples (Fig. 5A). Another line of evidence supporting the effectiveness of HHT treatment was that the asymmetry score in HHT-treated cells was biased towards the first half of ORFs (library L6, Fig. 1C).

**FIGURE 5:**
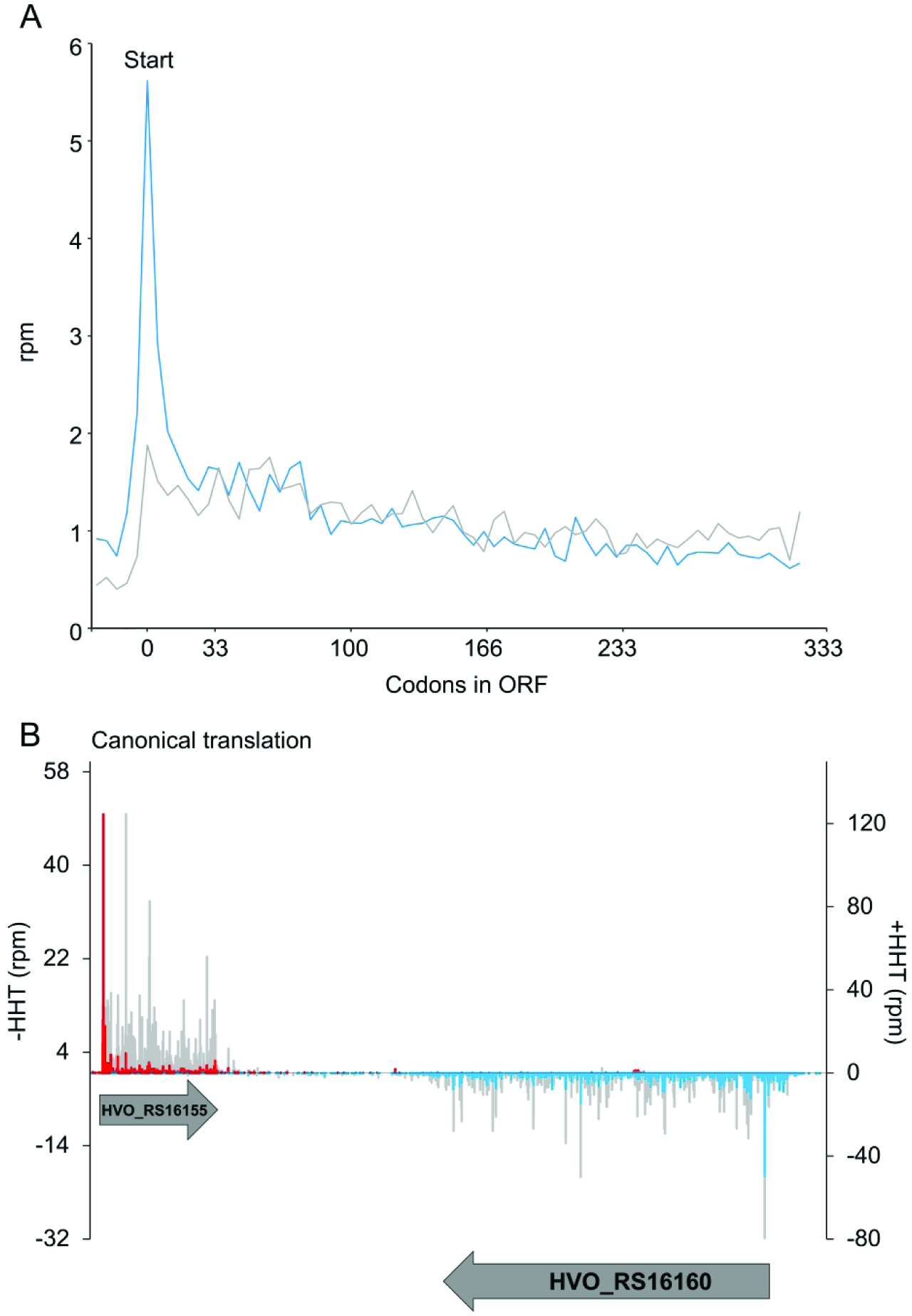
Harringtonine (HHT) locks ribosomes at translation start sites. (A) Meta-gene analysis plot of ribosome density along ORFs 1 kb downstream from the TSS. The blue line is HHT treatment and the gray line is no-drug treatment. (B) Representative plots of translation start site identification in annotated genes. Blue bars are HHT ribosome density on the minus strand, red bars are on the plus strand. Gray bars are ribosome density from untreated samples on both strands.

We then leveraged HHT-treatment to comprehensively identify initiation sites in *Hv* using previously established computational methods (26, 27). In brief, we identified all potential ORFs, asked if peaks of ribosome density aligned near the start codon of each potential ORF (± 3 nt), verified that these ORFs were translated in profiling data without HHT treatment (≥ 1 rpkm ribosome density), and assigned the type of TSS based on the gene annotation (NCBI RefSeq). Among the ORFs passing our analyses, 1,771 were previously annotated, representing 51% of annotated genes on the main chromosome. In several examples, the ribosome density on these genes was sharply increased at the start site in comparison to the rest of the ORF in HHT-treated but not in untreated samples (Fig. 5B). Yet there were other genes with little or no enrichment at the start codons. We cannot explain why some genes appear to be more sensitive to HHT than others. To increase the chances of finding bona fide start sites, we calculated the enrichment of ribosome density at the TSS for each potential ORF compared with the downstream density within the ORF. Candidates with high relative density (≥ 0.1) were considered HHT sensitive; 188 of annotated start sites were above this stringent threshold.

### Characterization of novel and alternative initiation

Using the same stringent parameters, we selected TSS that have not been annotated as start sites for further analysis. As described above, although a large proportion of annotated genes were validated through our approach, these annotated TSS only account for 92% of all of the identified TSS in the transcriptome, while the remaining 8% of TSS were novel (relative density ≥ 0.1). We classified the 160 remaining TSS as alternative TSS (aTSS) based on their relationship to the annotated ORF (Fig. 6A). We identified both small ORFs less than 50 codons (smORFs, 68, 43%) and novel unannotated TSS that encode large ORFs (18, 11%) in both intergenic regions and antisense to annotated genes. In addition, aTSS were identified upstream of annotated TSS (N-terminal extension, 31, 19%), and internal to annotated ORFs either in-frame (27, 17%) or out-of-frame (16, 10%). Many of the aTSS started with AUG (48%), although the known alternate start codons GUG (39%) and TTG (13%) were also observed (Fig. 6B).

**FIGURE 6:**
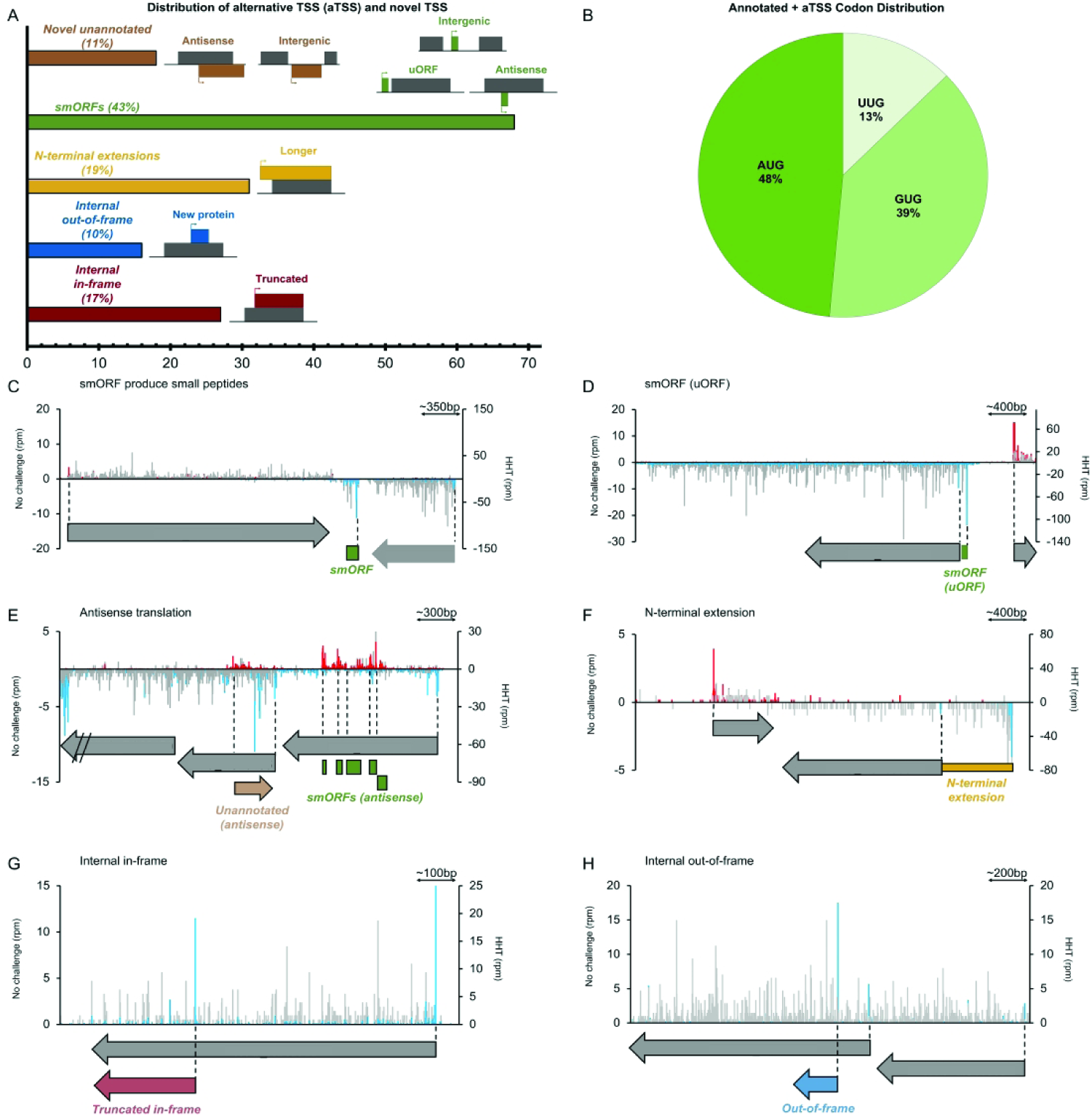
Identification of alternative translation initiation sites (aTSS). (A) Distribution of HHT-identified aTSS. (B) Distribution of initiation codons for HHT-identified aTSS. (C-H) Representative examples of aTSS identified in HHT-treated cells. (C) Intergenic smORF, (D) upstream smORF (uORF), (E) antisense translation initiation, (F) N-terminal extensions, (G) internal in-frame, and (H) internal out-of-frame initiation. Plus strand is plotted with positive values and minus strand with negative values. Untreated samples are shown with gray bars on both strands, HHT conditions on the plus strand are red bars, and blue bars are HHT conditions on the minus strand.

Small proteins have recently been found to be important players in a variety of processes in both Eukaryotes and Bacteria (27, 44–51) but have not been well characterized in Archaea. Using our ribosome profiling approach, we found many alternative TSS that were predicted to produce small proteins. smORFs were found in intergenic regions (Fig. 6C), as distinct proteins in 5’-UTRs of mRNAs as upstream ORFs (uORFs) (Fig. 6D), and antisense to coding regions in clusters (Fig. 6E). Many of these putative smORFs were highly conserved in other haloarchaea, suggesting their importance and functional potential.

We also observed N-terminal extensions of annotated genes, with enrichment of footprints at an upstream TSS in HHT-treated samples followed by elongating footprints extending to annotated ORF (Fig. 6F). In many cases peaks were also observed at the annotated TSS, suggesting that two distinct proteins may be produced from a single gene. For example, translation of a DEAD/DEAH box helicase likely initiated at its annotated TSS as well as an alternate TSS 273 nt upstream, encoding two isoforms of this helicase (Fig. 6F). This extended region was highly conserved at the amino acid level (86% identity, e-val = 2e-77).

Internal translational initiation could potentially produce N-terminal truncations (in-frame) or completely new proteins (out-of-frame) within a longer ORF, increasing the repertoire of proteins in a compact genome. One example is a gene encoding deoxyhypusine synthase that likely initiated at its annotated start site (ATG) and from an internal TSS 738 nt downstream in the same reading frame (Fig. 6G). The annotated deoxyhypusine synthase is a 359 amino acid long protein involved in post-translational modifications. The putative internal TSS potentially produces an N-terminally truncated protein only 113 amino acids long, losing the NAD/FAD-binding domain while retaining the transmembrane helix of deoxyhypusine synthase. In contrast, glutamate-5-semialdehyde dehydrogenase, involved in amino acid biosynthesis, initiated at the annotated TSS and also from an internal TSS 203 nt downstream in a different reading frame (Fig. 6H). This alternative frame could potentially produce a 75 amino acid protein as opposed to the annotated 444 amino acid protein. This novel internal out-of-frame protein was predicted to be highly disordered (89% of sequence) but had a significant domain hit (blastp, e-val = 2e-04) against an acyl transferase domain involved in polyketide synthesis.

Genes involved in the biosynthesis of amino acids and secondary metabolites (p=6.4E-4) and GTPase activity (p=3.7E-2) were enriched in the set of genes with N-terminal extensions, the latter including the cell division protein FtsZ and elongation factor 1-alpha. The internal TSS demonstrated a GO enrichment in universal stress proteins (UspA) and cell redox homeostasis (p=4.5E-2) for putative N-terminally truncated genes but no significant GO enrichment for out-of-frame TSS.

### Exploring translation of alternative TSS

Harringtonine treatment allowed us to use ribosome profiling to predict TSS at the global level, but the data could provide insight into the relative translational efficiencies of annotated and alternative TSS. We coupled standard ribosome profiling with RNA-seq to determine the translation efficiency for ORFs corresponding to the experimentally identified TSS in this study. Translation efficiency is a ratiometric estimation of the number of ribosomes per mRNA, where increases and decreases of efficiency are deviations from the baseline of 1:1 RPF and transcript levels. While the majority of aTSS and their corresponding genes were translated at levels comparable to their transcript levels, we observed outliers with high and low translation efficiency through a pair-wise comparison (Fig. 7). Because the 3’ UTRs of genes are not translated, their low translation efficiency values (Fig. 7, gray) could be used as a threshold to distinguish between poorly translated versus highly translated ORFs, a method established in Eukaryotes (42). Applying this type of analysis for newly identified ORFs from aTSS, we found that N-terminally extensions had more highly efficient translation compared to the other aTSS, even higher TE than their corresponding genes (Fig. 7, compare yellow and black), suggesting their functional importance.

**FIGURE 7:**
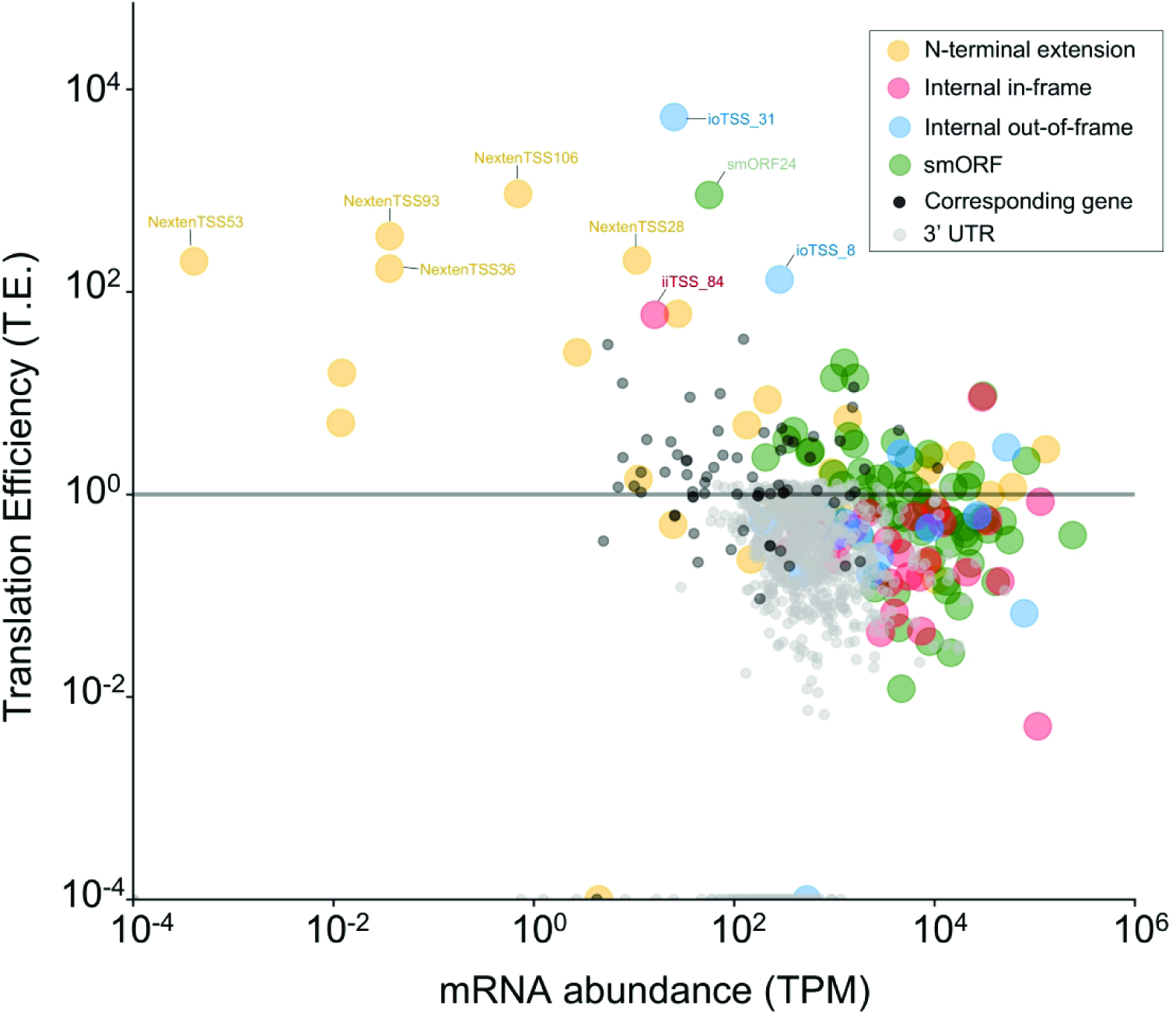
Translation of alternative TSS (aTSS). Scatterplot plot of translation efficiency of all discovered aTSS, their corresponding genes, and 3’ UTRs of these genes plotted against mRNA abundance. Translation efficiency was calculated from ribosome profiling and mRNA-seq data. N-terminal extensions are in yellow, smORFs in green, internal initiation sites that are in-frame in red, and internal out-of-frame initiation sites in blue. Corresponding genes are in black and 3’ UTRs are in gray, respectively.

## DISCUSSION

Translation is a highly regulated process and it represents the single largest investment of energy in the cell (52, 53). Studies of the mechanisms and regulation of protein synthesis in bacteria and eukarya have been greatly facilitated by the development of an experimental approach to globally analyze the full set of ribosomes engaged in translation, a high-throughput technique termed ribosome profiling. In comparison, due to a lack of comparable tools, we know relatively little about translation in Archaea (54, 55). Here we developed a ribosome profiling method in *Hv* and revealed the first global view of translation in an archaeon and in an extremophile. In the future, we anticipate that ribosome profiling will provide further insights into translational control of gene expression in *Hv* and that this protocol will be adapted to other members of the Archaea.

A critical challenge in the development of a robust and reproducible ribosome profiling protocol for a new organism is ensuring that translation is arrested in a way that faithfully captures the *in vivo* translational landscape. This problem is particularly acute when working with extremophiles because of the harsh experimental conditions required to maintain the integrity of their macromolecules; harvesting the culture places a strain on cells that can quickly and pervasively alter translation levels. In yeast, the preferred method is to harvest cultures by filtration followed by flash freezing; we were unable to obtain intact ribosomes from filtered *Hv* samples. Pre-treating the culture with a translation elongation inhibitor (anisomycin, ANS) inhibited translation well enough that we were able to obtain 70S monosomes from sucrose gradients after harvesting cultures by centrifugation. However, as reported for other microorganisms, pre-treatment with antibiotics led to problems with reproducibility, at least partially due to accumulation of ribosomes at the 5’-end of transcripts (given that initiation continues as elongation is arrested) (22, 30). To alleviate these issues, we used a direct flash-freezing method that allowed us to faithfully arrest translation after cell lysis and preserve the *in vivo* translational landscape. In this method, first reported for *E. coli* (22), cultures were directly flash-frozen with no antibiotics prior to cell lysis.

A second challenge in biochemical studies of protein synthesis in *Hv* is that the proteome and the ribosomes require high salt concentrations to maintain correct folding and intermolecular interactions (56). Previous studies in *Hv* have been unable to isolate intact 70S monosomes, probably due to subunit dissociation *in vitro* following cell lysis (57, 58). We were successful in developing lysis conditions and buffers that allow us to isolate 70S monosomes, a crucial step for ribosome profiling. Even with our optimized lysis buffer, however, we observed that monosome and polysome abundance declines over time after cell lysis, suggesting that ribosome subunits were dissociating with time. More optimization to resemble *Hv* intracellular salt concentrations (i.e. diversity of salts) may help. Still, the reproducibility we see in our profiling libraries indicates that this ribosome splitting is not biased towards any particular mRNA and does not affect the data overall. The challenge of ribosome instability *in vitro* will likely also affect attempts to perform ribosome profiling in other extremophiles such as hyperthermophiles and anaerobes, where environmental conditions are temperature and oxygen-limited. It may be that crosslinking strategies can stabilize 70S ribosomes on mRNA transcripts to overcome these challenges, although in our hands crosslinking did not noticeably alter monosome/polysome levels in *Hv* (data not shown).

Although we isolated a broad range of footprints (10 – 45 nt), we found that the major footprint from elongating ribosomes in *Hv* is 27 nt long. Because these footprints are trimmed back to the edges of the ribosome, they give the most precise information about the position of the ribosome. The fact that > 75% of 27 nt footprints map to the first position of codons indicates that ribosome profiling in *Hv* can capture reading frame at codon resolution, unlike studies in bacteria (22, 30). The size of the predominant *Hv* footprint is close to the size of the major eukaryotic footprint (28 nt in yeast), another example of the close evolutionary relationship between Archaea and Eukarya. This is surprising given that archaea have smaller ribosome subunits, more similar in size to bacterial than eukaryal ribosomes. However, the relatively higher number of translation factors found in archaea, most of which are homologous to eukaryotic translation factors, could conceivably produce a larger footprint (11). Alternatively, this larger footprint size may be related to the way rRNA pair with mRNA upstream of the P-site codon in *Hv*. Going forward it will be interesting to further characterize protein synthesis at the genome-wide level in other members of the third domain of life to shed light on this archaeal-eukaryal evolutionary relationship. In particular, studying translation and major footprint(s) of Asgard archaea, the closest evolutionary relative of the eukaryotic nucleus, will undoubtedly help towards this goal (8).

In addition to the predominant 27 nt footprints from elongating ribosomes, we also observed a distribution of shorter footprints (< 20 nt). In the context of a primarily leaderless transcriptome (>70% in *Hv*), ribosomes that initiate on a leaderless mRNA will have an mRNA channel that is empty upstream of the P site. After nuclease digestion, mRNA footprints from ribosomes on leaderless start codons will thus produce a smaller footprint size (peak at 16 nt) compared to an elongating ribosome (27 nt) that has the entire tunnel occupied by transcript (Fig. 3G). In contrast, ribosomes on start codons on leadered mRNAs have a longer footprint size due to the 5’ UTR sequence that occupies the mRNA channel during initiation. Indeed, we observed an enrichment of 27 nt footprints (enhanced with elongation inhibitors) upstream of start codons on the mRNAs that are leadered in *Hv.* This is of particular interest because our understanding of leaderless initiation, thought to be the evolutionarily oldest mechanism, is still poorly characterized (15, 55, 59). Our ability to distinguish between footprint sizes of leaderless and leadered transcripts, therefore, provides a model to address outstanding questions regarding the mechanisms between these different forms of initiation.

To our knowledge only two other studies have investigated the differences between leadered and leaderless mRNAs using ribosome profiling (60, 61). Neither reported differences in footprint lengths at start codons on leadered and leaderless transcripts but notable differences with their work and ours are that (i) the Bacteria studied did not have predominantly leaderless transcriptomes (*Mycobacterium smegmatis* 20%, *Streptomyces coelicolor* 21%), and (ii) a limited range of footprints sizes were size selected (*Mycobacterium smegmatis* 28 nt, *S. coelicolor* 26-32 nt) which may have prevented such analysis (60, 61). In the future, studies should isolate a broad distribution of footprint sizes to investigate how general footprint size differences are for leadered versus leaderless mRNAs across the three domains of life.

One of the strengths of ribosome profiling is its ability to detect differences in local elongation rate that occur as ribosomes pause during elongation. Ribosome pausing due to non-optimal codon usage or by interactions between the nascent peptide and the ribosomal exit tunnel can regulate gene expression (62). We used a targeted drug approach, treating *Hv* with serine hydroxamate (SHX), to starve the cells of a specific aminoacyl-tRNA and detect pauses at the level of codons. To our surprise, SHX did not yield pauses as Ser codons, as it does in bacteria, but instead caused strong pausing at Met codons. By measuring charged and uncharged tRNAs, we found evidence that SHX blocks the charging of Met-tRNA, either directly or by inhibiting Met biosynthesis (a serine-dependent process).

We further assessed the pausing landscape in *Hv*, using untreated cells, and found reproducible Pro pauses at A-, P-, and E-sites. Pro is known to be both a poor peptidyl acceptor and donor and polyproline sequences are particularly problematic. A specialized elongation factor (EFP in bacteria and eIF5A in eukaryotes) helps to mitigate these pauses by accelerating peptidyl transfer between Pro residues (63). Our data suggest that polyproline pausing also occurs in *Hv*, which encode a related elongation factor, aIF5A (64). Many of these pause effects were masked by ANS treatment, prior to harvesting (either flash freezing or centrifugation), likely because the drug inhibits elongation with some level of sequence selectivity. The observation of pauses reflecting known limitations in translation (e.g., Pro), and the disruption of these pauses by translation inhibitors, strongly suggests we are capturing true snapshots of *in vivo* biology that is not masked by methodology-based artifacts. Our results confirm that ribosome pausing occurs in Archaea with implications for gene regulation, an observation already demonstrated in the other domains of life (62, 65–70).

Mapping translation start sites (TSS) transcriptome-wide with antibiotics to trap initiation complexes has proven to be a powerful strategy in eukaryotes and bacteria (26, 27, 42). The need for accurate identification of TSS is particularly apparent in Archaea where most gene annotations are generated from general computational pipelines that are not totally reliable. Further complicating bioinformatics analyses, most genes identified in Archaea are not well conserved with known genes in other organisms. Using harringtonine (HHX) to lock ribosomes onto initiation sites, we accurately identified many TSS for annotated genes from the NCBI RefSeq annotation. We also identified potential protein-coding sequences in contexts that are often missed (71), such as TSS on extremely small ORFs or those that are within ORFs or antisense to known ORFs. We found hundreds of alternative TSS in *Hv* including initiation sites upstream of annotated TSS (N-terminal extensions), initiation sites within annotated ORFs that could produce truncated proteins (internal in-frame) or completely new proteins (internal out-of-frame), and small ORFs (<50 amino acids). These alternative proteins may have been previously obscured in proteomic data but may play important roles in cell physiology and stress response. N-terminally extended and internal initiation could play important roles in gene regulation of their corresponding ORFs by altering translation efficiency of the annotated protein, as proposed in Bacteria and Eukarya (26, 42). The verification of these putative alternative proteins is challenging in Archaea, compared to Bacteria and Eukarya, because genetic tools are less developed and the limitations on biochemical approaches due to proteomic adaptations to extreme environments (72).

## CONCLUSION

In conclusion, we coupled ribosome profiling with translation inhibitors to determine essential characteristics of translation in a member of the Archaea, for the first time. Specifically, we (i) determined the size of the archaeal ribosome footprint, (ii) assigned translation states of the ribosome to footprint lengths in a majority leaderless transcriptome, (iii) experimentally induced ribosome pauses and clarified the pausing landscape comprehensively, (iv) identified novel proteins including small open reading frames (smORFs), and (v) provided evidence that many genes initiate at alternative translation start sites (aTSS) around and within open reading frames (ORFs). This work demonstrates how a microorganism with a gene-dense genome can potentially produce proteins with distinct functions (isoforms) using the same gene. Lastly, ribosome profiling revealed the features archaea use in their translational apparatus, which are both a mosaic of bacteria and eukarya as well as features unique to their domain.

## AVAILABILITY

Data analysis scripts are available on github: https://github.com/dgelsin/Ribosome_profiling_MS

## ACCESSION NUMBERS

Ribosome profiling reads, density bigwigs, transcriptome reannotation, and translation start site annotations are available on NCBI GEO: GSE138990.

## ACKNOWLEDGEMENTS

We thank Dr. Rachel Green and her laboratory for reagents and helpful discussions. We thank Dr. Colin Wu for assistance in analyzing ribosome profiling data for Eukaryotes.

## FUNDING

This work was supported by the National Aeronautics and Space Administration [18-EXO18-0091 to J.D.], the Air Force Office of Scientific Research [FA9950-14-1-0118 to J.D.], and the National Institutes of Health [GM110113 to A.B.]. Funding for open access charge: National Aeronautics and Space Administration.

## SUPPLEMENTAL FIGURE LEGENDS

**FIGURE S1:**
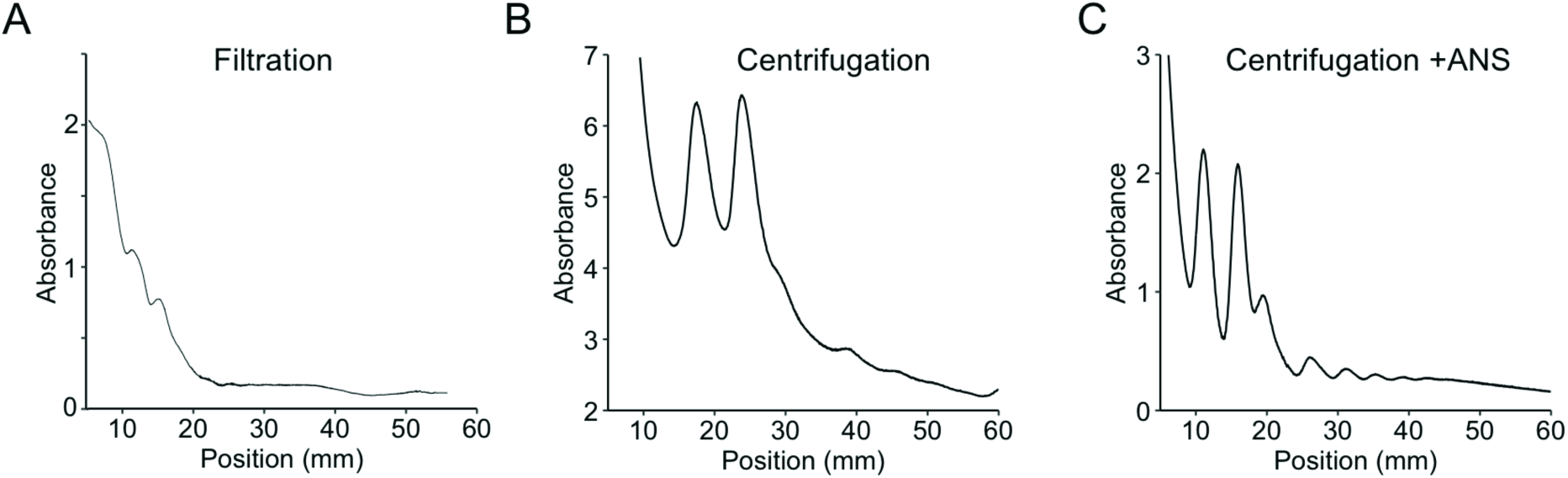
Sucrose gradient plots of (A) cells filtered and harvested with liquid nitrogen, (B) cells harvested with centrifugation and no-drug added, and (C) cells harvested with centrifugation with anisomycin (ANS) added.

**FIGURE S2:**
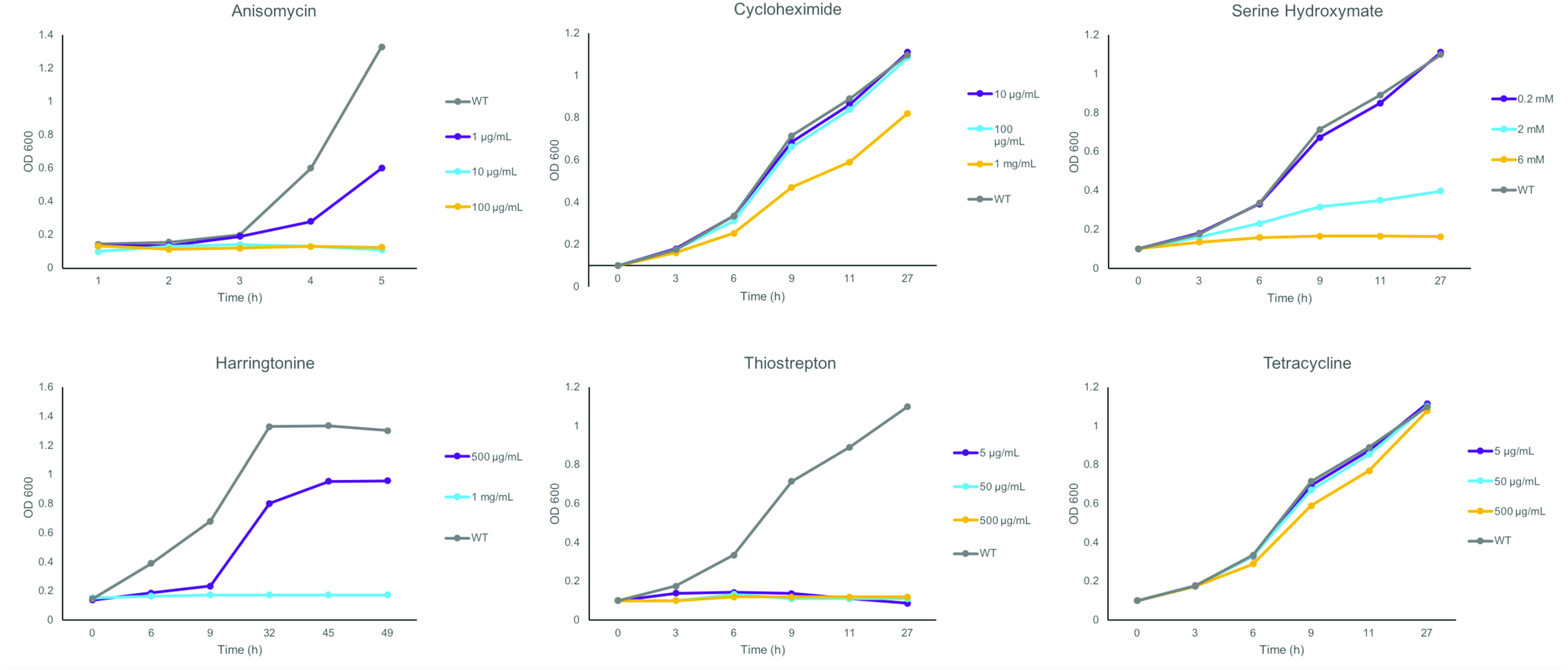
Growth curves of *Hv* exposed to translation inhibitors.

**FIGURE S3:**
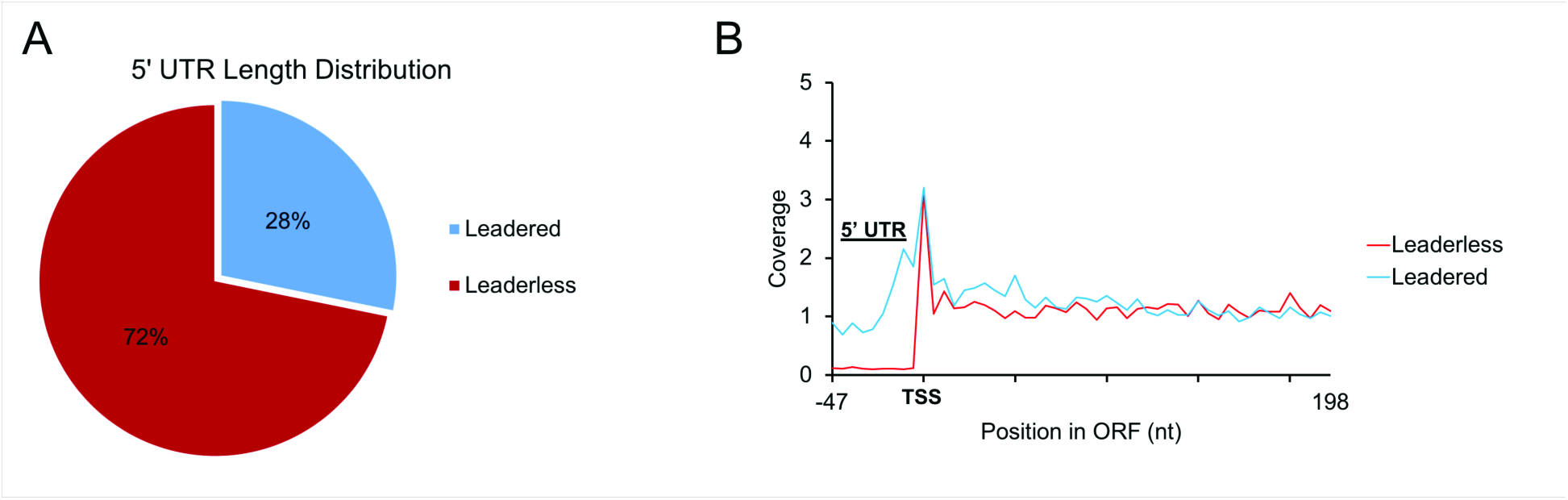
Dependency of ribosome integrity in *Hv* on potassium chloride (KCl) levels shown with sucrose gradient plots.

**FIGURE S4:**
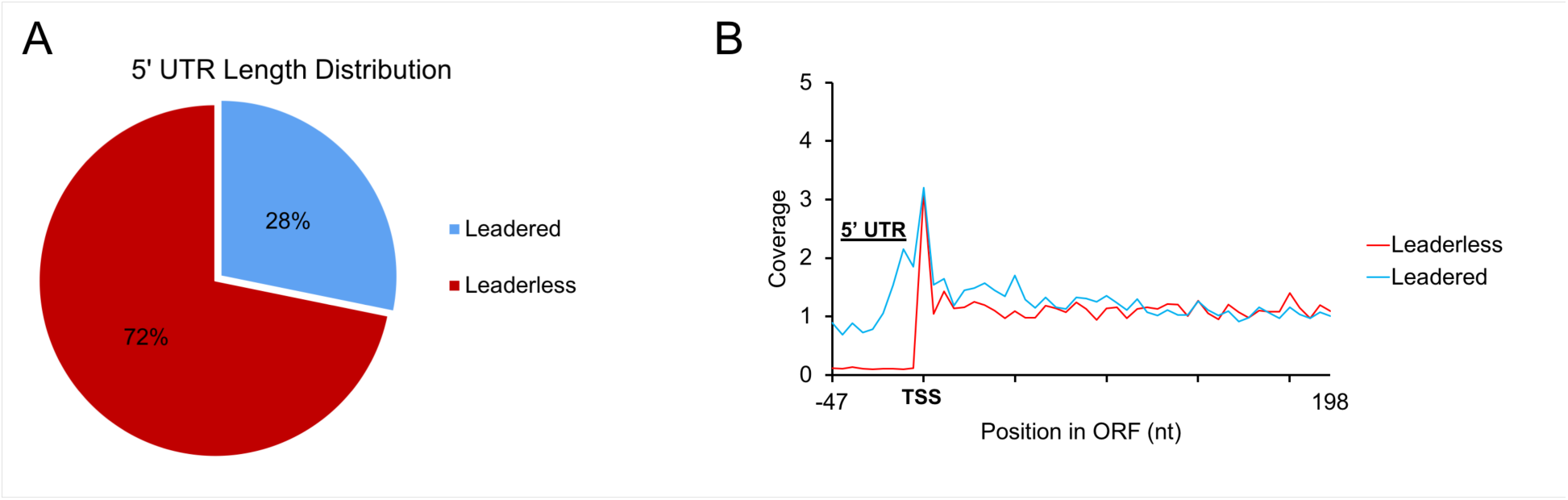
Experimental identification of leadered and leaderless mRNAs in *Hv.* (A) 5’ UTR distribution of the transcriptome in *Hv.* (B) Meta-gene analysis plot of RNA-seq reads on leadered and leaderless transcripts.

**FIGURE S5:**
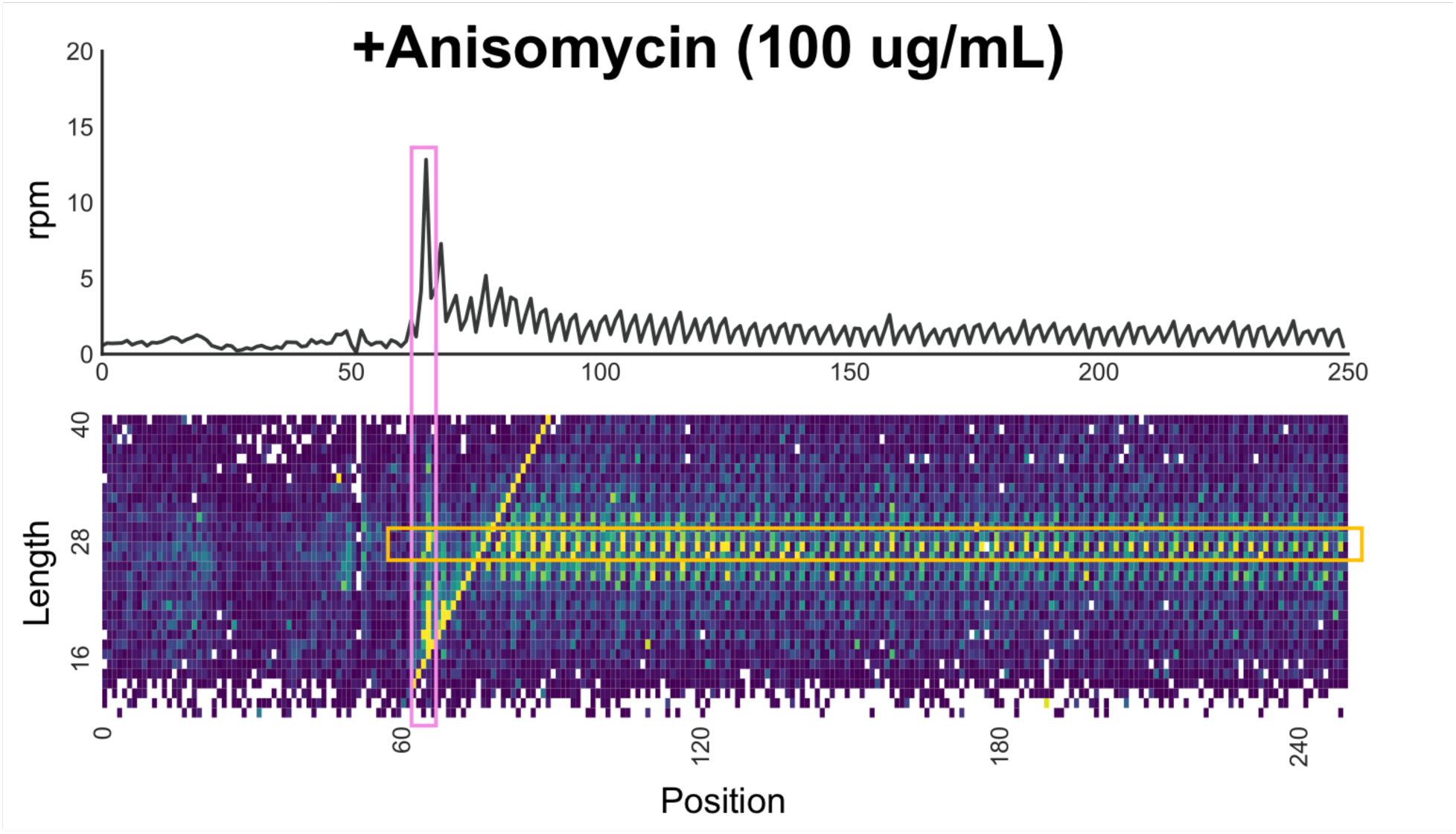
3’-mapped meta-gene analysis of all footprints (10–40) for (A) no drug and (B) anisomycin-treated cells.

## Notes

https://www.ncbi.nlm.nih.gov/geo/query/acc.cgi?acc=GSE138990

